# Cholesterol-mediated tight junction formation

**DOI:** 10.1101/2022.06.30.498218

**Authors:** Kenta Shigetomi, Junichi Ikenouchi

## Abstract

Lipids have the ability to self-organize but the significance of this phenomenon in the assembly of membrane structures is unknown. We previously reported that cholesterol enrichment directs the formation of tight junctions (TJs), adhesion structures responsible for the epithelial barrier (Shigetomi et al., 2018). However, it is unclear how cholesterol accumulates and informs TJ formation. Cholesterol typically accumulates in the vicinity of apical cell junctions (Shigetomi et al., 2018). Surprisingly, cholesterol distribution is unaltered in an epithelial cell line that lacks TJs, generated by suppressing the expression of claudins, membrane proteins that determine the barrier properties of TJs. Assembly of claudin into hallmark ‘strands’ is canonically thought to require binding to Zonula occludens (ZO) proteins but a claudin mutant that cannot bind to ZO proteins still form strands. These results suggest a novel mechanism for TJ formation that is dependent on cholesterol and reveal an unexpected role of ZO proteins as organizers of the cholesterol domain.

## Introduction

Epithelial cells adhere to each other to form a continuous cell sheet, which covers the surface of organs. The epithelial cell sheet also functions as a barrier that separates the inside of the body from the outside and limits free diffusion of small substances such as ions. This function is assumed by TJs that are located at the most apical side of the lateral membrane. Claudins, four transmembrane proteins, and Zonula occludens (ZO) proteins, scaffolding proteins for claudins, are the major constituents of TJs (Tsukita et al., 2001). The widely accepted text book model of TJ formation proposes that incorporation of claudins into the characteristic sealing strands is regulated by the scaffolding function of ZO proteins. In other words, multimerization of ZO proteins at the apical junctional complexes (AJCs) serves as a platform to enable claudin polymerization (Beutel et al., 2019; Umeda et al., 2006).

Confoundingly, our previous study revealed that recruitment of ZO proteins to AJCs is insufficient to induce TJ formation. Tight junctions are membrane domains rich in cholesterol and sphingomyelin (SM) with very long chain fatty acids (Nusrat et al., 2000; Shigetomi et al., 2018). Removal of plasma membrane cholesterol by treatment with methyl-beta-cyclodextrin (MbCD) promptly disrupts TJs by inducing rapid removal of claudins from the plasma membrane without disturbing the localization of ZO proteins, which suggests that cholesterol-rich membrane domains, rather than ZO proteins themselves, informs TJ formation(Shigetomi et al., 2018). However, it is unclear that the cholesterol-rich membrane domain is necessary and sufficient to induce TJs; it also remains possible that the presence of polymerized claudins enables the membrane domain to form. Furthermore, since the formation of AJs was not affected by MbCD treatment (Shigetomi et al., 2018), the formation mechanisms of AJ and TJ clearly differ in terms of the requirement for cholesterol. Therefore, in order to specifically address the mechanism of cholesterol accumulation at TJs and the reason why TJ formation depends on cholesterol, we established epithelial cells in which all claudin isoforms were knocked out by CRISPR.

## Results and Discussion

### Establishment of epithelial cells lacking TJs by knock-out of all expressed claudin isoforms

We generated claudin-null cells lacking all expressed claudin isoforms from the mouse cultured mammary gland epithelial cell EpH4 (claudin-1, −3, −4, −7, −9, and −10) (Yamazaki et al., 2011) by CRISPR-Cpf1-mediated gene knockout. The loss of claudin isoforms’ expression in claudin-null cells was confirmed by immunofluorescence microscopy and western blotting (Figs. 1A and 1B). Freeze-fracture replica electron microscopy revealed that TJ strands, typically observed beneath microvilli in wild-type EpH4 cells, were never observed in claudin-null cells (Fig. 1C). We also examined the localization of other TJ proteins in claudin-null cells. One of the ZO proteins, ZO-1, was enriched at AJCs in claudin-null cells but their amount increased relative to wild-type (Fig. 1D). Like claudins, occludin is a TJ membrane protein that is also recruited to AJCs via ZO proteins and its amount proportionately increased (Fig. 1D). In addition, the accumulation of JAM-A also increased in claudin-null cells (Fig. 1D).

**Figure 1.**
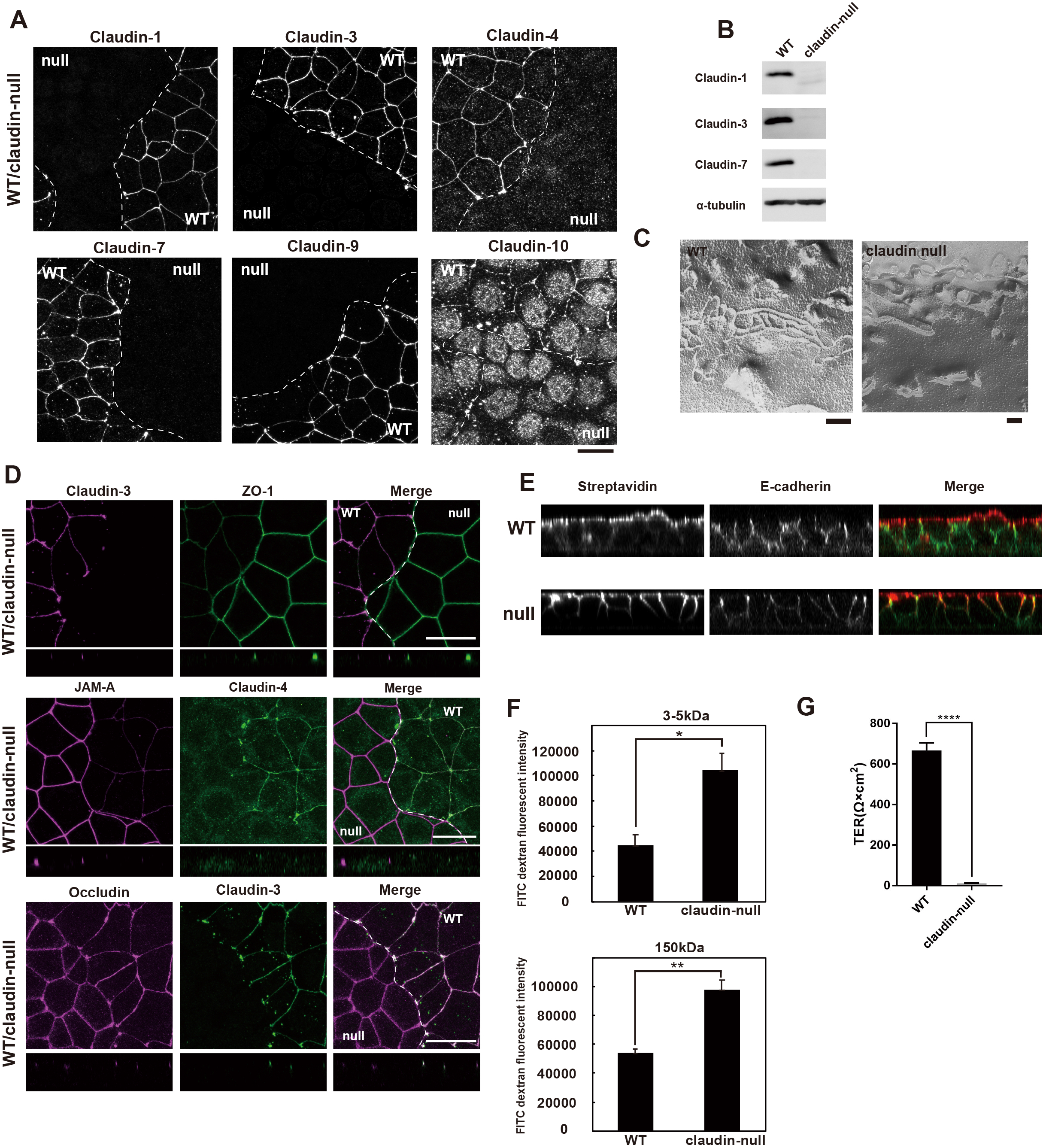
Establishment of claudin-null cells. (A) Representative immunofluorescence images of a co-culture of EpH4 WT and claudin-null cells stained for claudin-1, claudin-3, claudin-4, claudin-7, claudin-9 and claudin-10. Scale bar, 20 µm. (B) Immunoblotting of whole cell lysates of EpH4 WT and claudin-null cells. Claudin-1, claudin-3 and claudin-7 were not detected in claudin-null cells. (C) Freeze-fracture EM images of TJ strands in EpH4 WT and claudin-null cells. TJ strands were never observed in claudin-null cells. Scale bar, 200 nm. (D) Representative immunofluorescence images of a co-culture of EpH4 WT and claudin-null cells stained for claudin-3 and ZO-1 (upper panel), JAM-A and claudin-4 (middle panel) or occludin and claudin-3 (lower panel). Scale bar, 20 µm. (E) Sulfo-NHS-biotin was added to the apical compartment to visualize the paracellular tracer flux. The distribution of biotin was detected by staining of streptavidin. Lateral membranes are marked by E-cadherin. Note that in claudin-null cells, the biotin appears free to take the paracellular route over TJs. (F) Apical-to-basal permeability of 3-5-kD and 150kD FITC-dextran was significantly increased in claudin-null cells. (G) Trans-epithelial resistance (TER) was significantly decreased in claudin-null cells indicating loss of epithelial barrier in claudin-null cells. (Student’s t test, * p,0.05, ** p<0.01, **** p<0.001)

Based on our observations so far, it should follow that the barrier function of claudin-null cells is impaired. Indeed, in the biotin tracer assay, only the apical membrane is biotinylated in wild-type cells, as visualized by FITC-conjugated streptavidin. In contrast, biotinylation is detected at the lateral membrane in claudin-null cells, indicating that the biotin tracer freely passed through the intercellular spaces (Fig. 1E). We also tested the barrier function against macromolecules in claudin-null cells using FITC-dextran with molecular weights of 3 to 5 kDa and 150 kDa (Fig. 1F). In claudin-null cells, the permeability of both molecular weight macromolecules was significantly increased as compared to wild-type cells. Finally, we confirmed the loss of the TJ barrier in claudin-null cells by transepithelial electric resistance (TER) measurement (Fig. 1G). Therefore, we concluded that TJs are structurally and functionally absent in claudin-null cells.

### Claudin null cells are normally polarized but junctional contractility is aberrantly up-regulated

As TJs are thought to function as a fence at the boundary between apical and basolateral membranes, which contributes to epithelial polarization, we next examined the apicobasal polarity of claudin-null cells. When the apical marker proteins GFP-tagged podocalyxin-1 and prominin-2 were expressed in claudin-null cells, both localized at the apical membrane as well as in wild-type cells (Fig. 2A). Na-K ATPase α1, a lateral membrane marker protein, was correctly localized at the lateral membrane in claudin-null cells (Fig. 2B). Furthermore, Par-3, an essential component of the polarity protein complex, was concentrated at AJCs in claudin-null cells as in wild-type cells (Fig. 2C). These results indicate that cell polarity is maintained even in the absence of TJs, consistent with previous studies (Ikenouchi et al., 2012; Otani et al., 2019; Umeda et al., 2006).

**Figure 2.**
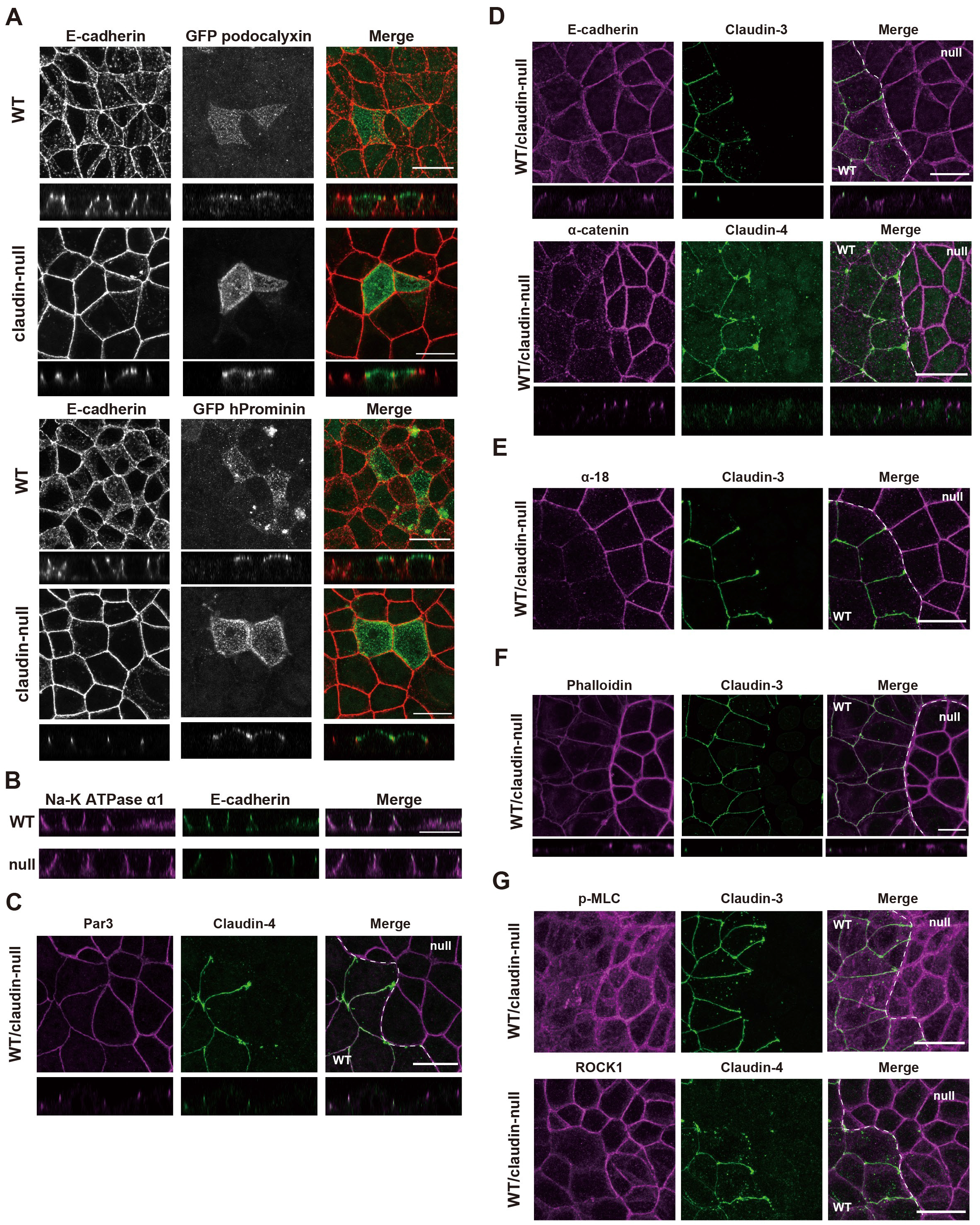
Apico-basal polarity is normally maintained in claudin-null cells. (A) GFP-tagged podocalyxin and GFP-tagged prominin2 were expressed in EpH4 WT and claudin-null cells. GFP podocalyxin and prominin2 localized to apical membrane in both EpH4 WT and claudin-null cells. Scale bar, 20 μm (B) Representative immunofluorescence images of a co-culture of EpH4 WT and claudin-null cells stained for Na-K ATPase. Scale bar, 20 μm. (C) Representative immunofluorescence images of a co-culture of EpH4 WT and claudin-null cells stained for Par3. Scale bar, 20 μm. (D) Representative immunofluorescence images of a co-culture of EpH4 WT and claudin-null cells stained for E-cadherin and α-catenin. Scale bar, 20 μm. (E) Representative immunofluorescence images of a co-culture of EpH4 WT and claudin-null cells stained with α18 antibody, which recognizes open-form α-catenin specifically. Scale bar, 20 μm. (F) Representative immunofluorescence images of a co-culture of EpH4 WT and claudin-null cells stained with phalloidin. Scale bar, 20 μm. (G) Representative immunofluorescence images of co-culture of EpH4 WT and claudin-null cells stained for phosphorylated myosin light chain (pMLC) and ROCK1. Scale bar, 20 μm.

Next, we investigated the localization of proteins associated with adherens junctions (AJs). In claudin-null cells, E-cadherin and α-catenin were concentrated more at AJCs than in wild-type cells (Fig. 2D). α-Catenin undergoes a tension-dependent conformational change. We noted a marked increase in the open form of α-catenin at AJCs as determined by staining with the α-18 antibody that recognizes only the open form of α-catenin, suggesting that the intercellular tension is increased in claudin-null cells (Fig. 2E). Accordingly, the circumferential actin ring underlying AJs was more robust (Fig. 2F) and myosin activity (p-MLC) was elevated in association with increased ROCK accumulation at AJs in claudin-null cells (Fig. 2G). Thus, the contractile force of the circumferential actin belt was increased, indicating that Rho activity at AJCs is up-regulated in claudin-null cells. Recent reports suggest that when TJs partially collapse, transient, localized Rho activation, or ‘Rho flares’, occurs at AJCs to repair the barrier function of TJs by ROCK-mediated contraction of AJCs (Stephenson et al., 2019; Varadarajan et al., 2019). Thus, we may be observing a similar response to the loss of TJs barrier in claudin-null cells.

### Polymerization of claudins to TJ strands does not require interaction with ZO proteins

Importantly, exogenous expression of GFP-tagged wild-type claudin-3 (WT; rescue cells are WTres) rescued the various phenotypes of claudin-null cells. Claudin-3 co-localized with ZO-1 and reversed its aberrant accumulation at AJCs (Fig. 3A). Freeze-fracture electron microscopy revealed that WTres cells have well-developed TJ strands (Fig. 3B). TER measurements revealed that functional TJs were also recovered in WTres cells (Fig. 3C). The staining intensities of α18, phalloidin, ROCK, and p-MLC were attenuated in WTres cells, suggesting that expression of WT claudin-3 restored the TJ barrier and reduced the contractility at AJCs (Fig. 3D).

**Figure 3.**
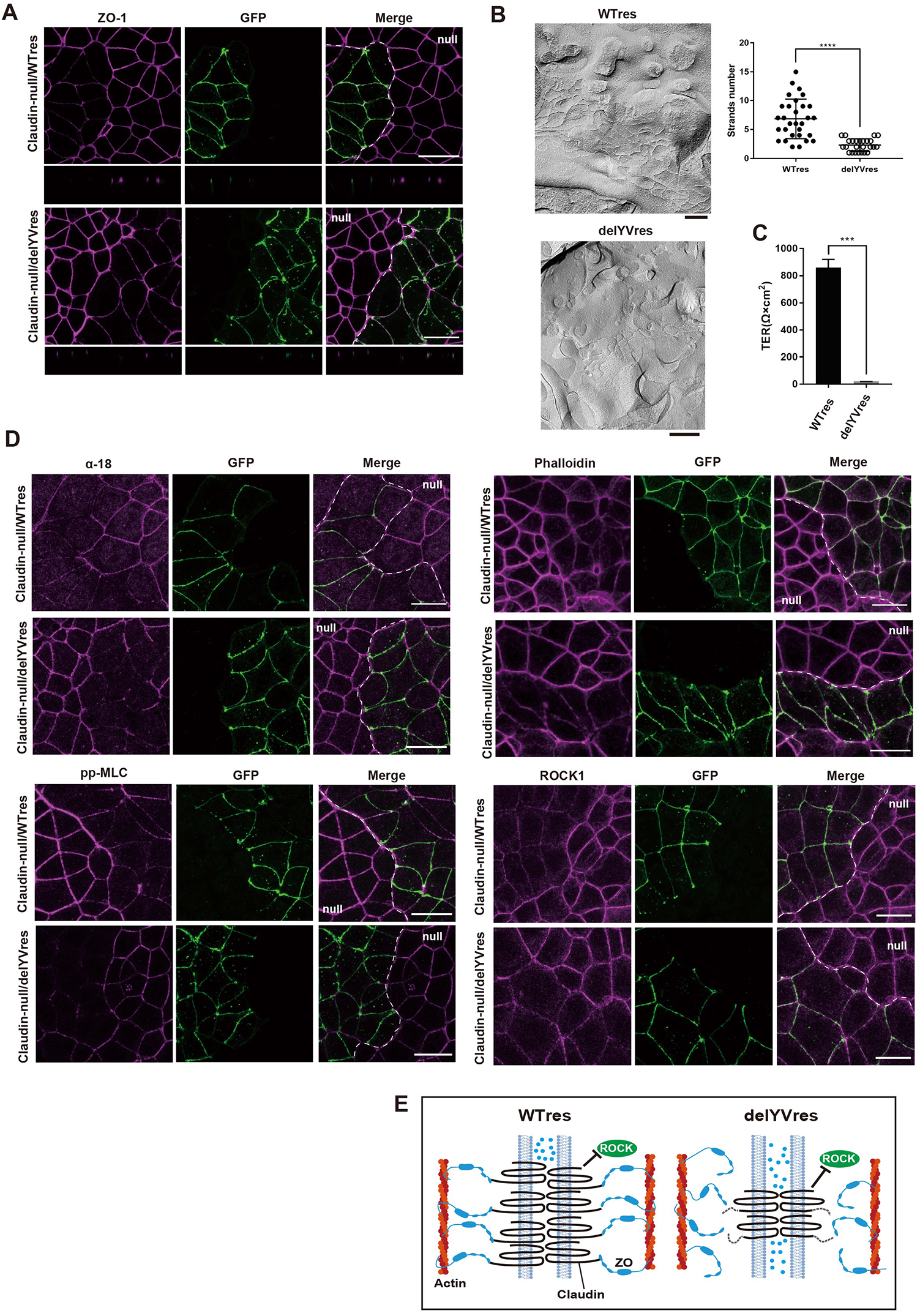
Claudin-3 delYV mutants induce TJ strand formation in claudin-null cells independently of ZO proteins. (A) Representative immunofluorescence images of a co-culture of claudin-null and WT res or delYV res cells stained for ZO-1. Scale bar, 20 μm. (B) Freeze-fracture EM images of TJ strands in WTres and delYVres cells. Number of TJ strands were quantified based on the freeze-fracture EM images. Scale bar, 200 nm. (C) TER was significantly decreased in delYVres cells as compared to WTres cells (Student’s t test, *** p<0.001). (D) Representative immunofluorescence images of a co-culture of claudin-null and WT res or delYV res cells stained for open-form α-catenin, phosphorylated myosin light chain, actin cytoskeleton and ROCK1. Scale bar, 20 μm. (E) Schematic diagram about the relationship between ZO proteins and claudin-3 WT or delYV mutant.

The C-terminal PDZ binding motif of claudin is well conserved among claudin isoforms and binds to the PDZ domain of ZO proteins. It is long assumed that the binding of claudin to ZO proteins is essential for TJ formation since TJs are not formed in cells lacking ZO proteins (Fanning et al., 2012; Umeda et al., 2006). However, this assumption was never directly tested in previous studies. Therefore, we expressed a mutant claudin-3 lacking the PDZ binding motif (claudin-3 delYV) in claudin-null cells (delYVres) to examine whether claudin-ZO interaction is required for TJ formation. To our surprise, claudin-3 delYV, which cannot bind to ZO proteins, was enriched at AJCs where it co-localized with ZO-1 (Fig. 3A). This result indicates that binding to ZO proteins is unnecessary for claudins to accumulate at AJCs. Remarkably, freeze-fracture electron microscopic analyses revealed bona fide TJ strands in delYVres cells, indicating that claudins are able to polymerize in the absence of ZO proteins (Fig. 3B). Nevertheless, the number of TJ strands was significantly decreased in delYVres cells compared to WTres cells (Fig. 3B). Therefore, it was unsurprising that TER was undeveloped (Fig. 3C) and that the paracellular barrier was severely compromised (Fig. 5D) in delYVres cells. By contrast, the up-regulation of actomyosin contractility associated with AJCs in claudin-null cells was cancelled in delYVres cells (Fig. 3D). Thus, actomyosin contractility is enhanced in claudin-null cells by a mechanism distinct from the ‘Rho flares’ described previously since barrier function and actomyosin contractility appear to be regulated independently (Fig. 3E). It is unclear currently how the formation of TJ strands regulates contractility of actomyosin but it is an interesting research topic in the future.

### Cholesterol is highly accumulated at the outer leaflet of the plasma membrane at AJCs

The unexpected finding that the claudin-3 delYV mutant accumulates in the vicinity of AJCs and polymerizes to form TJ strands despite its inability to associate with ZO proteins suggests that these processes can proceed in a ZO proteins-independent manner. The plasma membrane of AJCs is relatively enriched in cholesterol and SM with very long chain fatty acids (Fig. 4A) (Shigetomi et al., 2018). Moreover, cholesterol removal from the plasma membrane with MbCD impairs claudin accumulation without altering the localization of ZO proteins (Shigetomi et al., 2018). Therefore, we hypothesized that the formation of TJ strands in delYVres cells is driven by the accumulation of cholesterol at AJCs.

**Figure 4.**
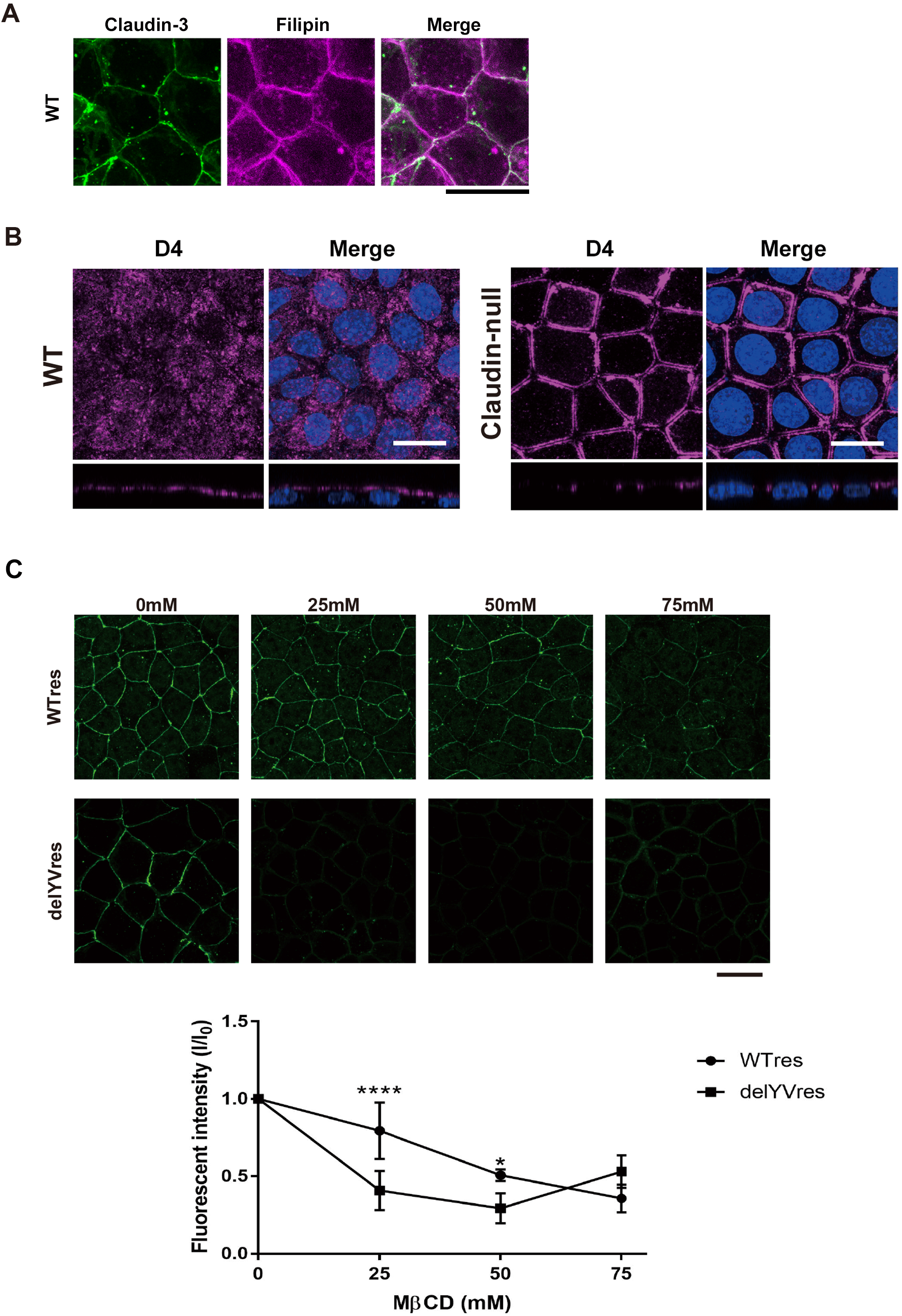
Accumulation of cholesterol at AJCs is maintained without TJs in claudin-null cells. (A) Distribution of cholesterol visualized by filipin staining in EpH4 cells expressing GFP-claudin3. Scale bar, 20 µm. (B) Distribution of cholesterol at the outer leaflet of plasma membrane visualized by D4 staining in EpH4 WT and claudin-null cells. Scale bar, 20 µm. (C) Changes of subcellular localization of GFP-claudin3 WT and GFP-claudin3 delYV when the WTres and delYVres cells were treated with MbCD at various concentration. Scale bar, 20 µm.

The protein-based probe derived from domain 4 of perfringolysin O (D4) derived from *Clostridium perfringens* specifically binds to cholesterol (Ishitsuka et al., 2004; Shimada et al., 2002; Yamaji et al., 1998). Due to its molecular size, D4 should stain the apical membrane only, and not stain the lateral membrane, as long as the TJ barrier is intact. As expected, D4 stained the apical membrane in wild-type cells (Fig. 4B). Strikingly, cholesterol was highly concentrated at the apical region of the lateral membrane where we would expect AJCs (Fig. 4B). This was notable as the barrier against FITC-dextran 150 kDa was lost in claudin-null cells (Fig. 1F). That D4 only recognized the outer leaflet of a limited apical region of the lateral membrane clearly indicates that the accumulation of cholesterol at AJCs is maintained in claudin-null cells.

Considering that TJ strands form independently of claudin binding to ZO proteins and that cholesterol is still enriched near AJCs in the absence of claudin assembly, we supposed that claudins concentrate at AJCs through their affinity for cholesterol-rich membrane domains and thereafter self-assemble into TJ strands. If this were true, claudin-3 delYV mutant would be more sensitive to cholesterol removal than WT claudin-3 since the former lacks the anchor to the circumferential actin ring through ZO proteins. Accumulation of WT claudin-3 at AJCs was unaffected by treatment with up to 50 mM MbCD. By contrast, 25 mM MbCD was enough to weaken the accumulation of mutant claudin-3 at AJCs (Fig. 4C). These results clearly support an active role for cholesterol-rich membrane domains in concentrating claudin at AJCs.

### Palmitoylation mediates claudin association with the cholesterol-rich membrane domain at AJCs

Then, what determines the affinity of claudin for such membrane domains? Palmitoylation is an important post-translational modification that determines the localization of membrane proteins at the plasma membrane (Diaz-Rohrer et al., 2014). Molecular dynamics simulation in a recent study predicts that claudin palmitoylation is important for its localization at the plasma membrane through the interaction with cholesterol and SM (Rajagopal et al., 2019). In fact, four cysteine residues of claudin-14 reportedly undergo palmitoylation and these palmitoylation sites are well-conserved among claudin isoforms (Van Itallie et al., 2005). Therefore, we next considered whether palmitoylation is involved in claudin targeting to AJCs by generating a claudin-3 mutant in which the conserved membrane-proximal cysteines were mutated to serine (4S mutant) (Fig. 5A). Immunofluorescence examination of the stably-expressed claudin-3 4S mutant in the claudin-null background (4Sres cells) showed the 4S mutant to localize uniformly to the lateral membrane with virtually no enrichment at AJCs (Fig. 5B). In addition, TJ strands were never observed in 4Sres cells (Fig. 5C). We also confirmed that barrier function was not restored in 4Sres cells (Fig. 5D). Furthermore, contractility of the circumferential actin ring was still up-regulated in 4Sres cells similar to claudin-null cells, leading us to conclude that claudin palmitoylation is necessary for TJ formation (Fig. 5E). This was in stark contrast to the localization of D4, which unwaveringly stained the apical membrane around AJCs (Fig. 5F). Therefore, we concluded that cholesterol accumulation at AJCs is a requisite precondition to TJ formation by informing claudin enrichment through its palmitoylated moiety.

**Figure 5.**
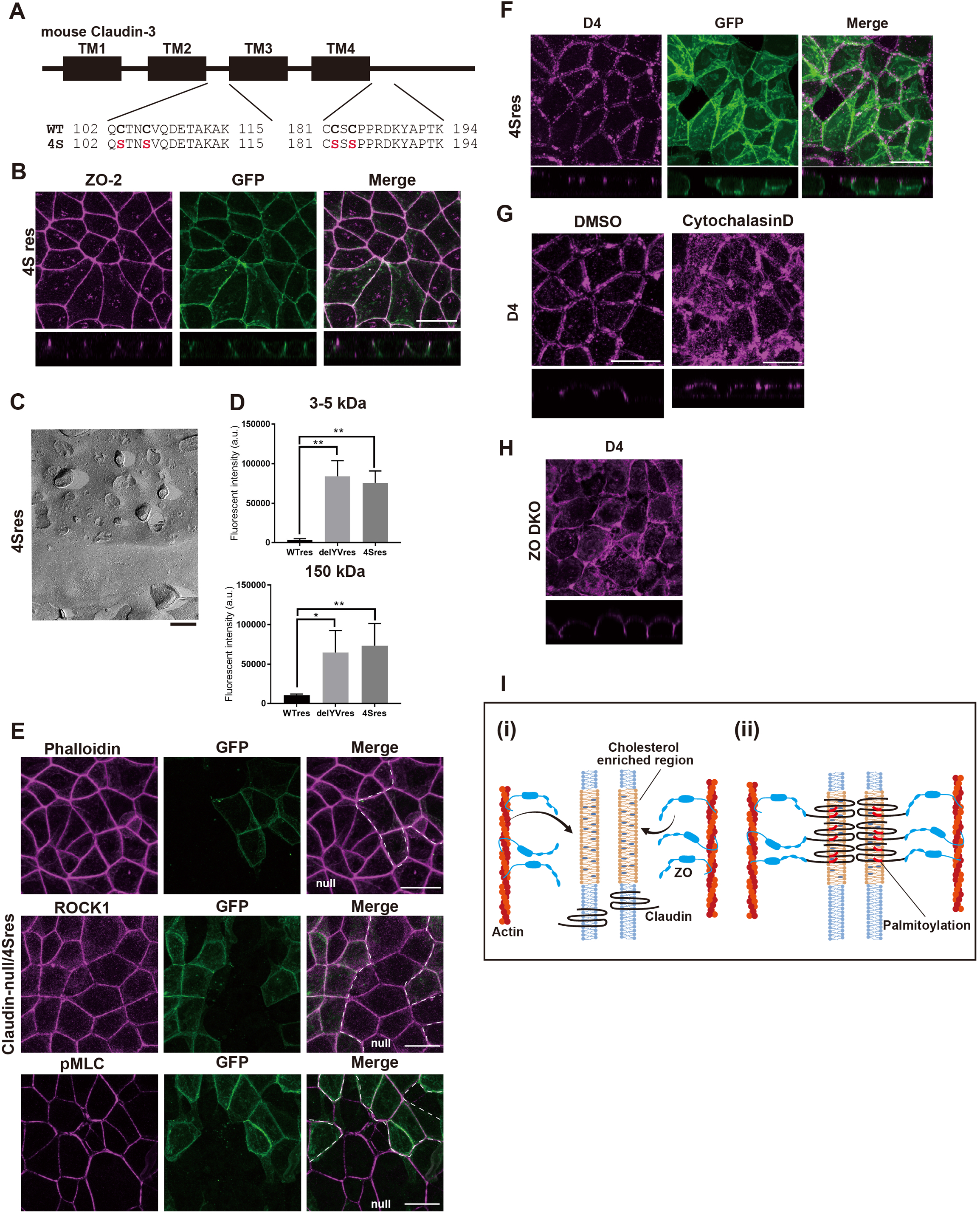
Palmitoylation of claudin required for TJ formation. (A) Schematic diagram of claudin-3 to illustrate its four transmembrane domains (TM1-TM4). Sequences around the membrane-proximal cysteines are shown. The cysteine-to-serine mutations are shown below the wild-type sequence. (B) Representative immunofluorescence images of 4Sres cells stained for ZO-2. Scale bar, 20 μm. (C) Freeze-fracture EM images of TJ strands in 4Sres cells. Scale bar, 200 nm. (D) Apical-to-basal permeability of 3-5-kD and 150kD FITC-dextran was compromised in delYVres and 4Sres cells. (E) Representative immunofluorescence images of a co-culture of claudin-null and 4Sres cells stained for phosphorylated myosin light chain, actin cytoskeleton and ROCK1. Scale bar, 20 μm. (F) Distribution of cholesterol at the outer leaflet of plasma membrane visualized by D4 staining in 4Sres cells. Scale bar, 20 µm. (G) Distribution of cholesterol at the outer leaflet of plasma membrane visualized by D4 staining in claudin-null cells treated with 1 μM cytochalasinD. Scale bar, 20 µm. (H) Working model of how claudins accumulate to AJC and form TJs independently of binding to ZO proteins. (i) Circumferential actin ring directs accumulation of cholesterol in the vicinity of AJC. ZO proteins possibly mediate the enrichment of cholesterol directly or indirectly. (ii) Palmitoylated claudins are recruited to the cholesterol-rich membrane domains, resulting in the assembly of claudins and TJ formation claudins. (Two-way ANOVA, * p,0.05, ** p<0.01, **** p<0.001)

So far, we have demonstrated that the accumulation of cholesterol at AJCs is a key determinant of claudin localization in epithelial cells but how does cholesterol accumulate at AJCs? Although not a precise parallel, the circumferential actin ring at AJCs is similar to the contractile ring formed during cytokinesis in that both are contractile actomyosin ring structures associated with the plasma membrane. Intriguingly, cholesterol accumulates at the cleavage furrow during cytokinesis in an actin-dependent manner as disruption of the contractile actin ring by cytochalasin D treatment impairs cholesterol accumulation at the cleavage furrow (Abe et al., 2012; Ng et al., 2005). Therefore, we examined the effects of cytochalasin D treatment on cholesterol distribution in claudin-null cells. We found that cholesterol accumulation at AJCs was abolished and it was instead diffusely localized throughout the entire apical membrane, indicating that an analogous mechanism to the cleavage furrow is at work here (Fig. 5G).

Previous studies suggest that the formation of the contractile ring and the enrichment of cholesterol at the cleavage furrow are interdependent (Abe et al., 2012; Ng et al., 2005). Inhibiting cholesterol accumulation at the cleavage furrow causes cytokinesis failure by impairing contractile ring formation. Therefore, we tested whether the formation of the circumferential actin ring is similarly dependent on cholesterol accumulation at AJCs. Cholesterol removal by MbCD treatment had no effect on circumferential actin ring formation in claudin-null cells, leading us to conclude that the circumferential actin ring preceded cholesterol accumulation at AJCs and not vice versa (Fig. S1).

When we previously reported that epithelial cells lacking ZO-1 and ZO-2 (ZO dKO cells) fail to form TJs, our interpretation of this result was that ZO proteins, as claudin scaffolds, determine where TJ strands are polymerized (Ikenouchi et al., 2007; Umeda et al., 2006). However, the delYV mutant lacking the association with ZO proteins still accumulates at AJCs and forms TJ strands, implying that binding to scaffolding proteins is dispensable for claudin accumulation. Therefore, we re-examined the role of ZO proteins in TJ formation by regarding them as membrane domain organizers instead. Consistent with this viewpoint, cholesterol was no longer concentrated at AJCs but broadly distributed to the lateral membrane (Fig. 5H).

The crux of this study is the observation that the circumferential actin ring underlying AJCs causes the formation of a cholesterol-rich membrane domain, which is essential for the formation of TJs. Altogether, our results strongly argue for a reconsideration of TJ formation, whereby protein-lipid (claudin-cholesterol) interaction—not protein-protein (claudin-ZO proteins) interaction as canonically believed—informs claudin strand polymerization. (Fig. 5I). Elucidating how the formation of cholesterol-rich domains at AJCs is induced by ZO proteins and the circumferential actin rings is an important research topic for the future. Relevant to this question, it was recently reported that cholesterol-rich liquid order domains are increased in T cells by artificially stabilizing the actin cytoskeleton (Dinic et al., 2013). There are several possibilities as to how non-labile actin structures could cause the accumulation of cholesterol and SM to the outer leaflet of the overlaying plasma membrane. It has been suggested that the actin cytoskeleton immobilizes phosphatidylserine (PS) with long-chain saturated fatty acids, resulting in the formation of membrane domains containing phospholipids with saturated fatty acids, including sphingomyelin, GPI-anchored protein and cholesterol on the outer leaflet by trans-bilayer coupling (Raghupathy et al., 2015). Such trans-bilayer coupling of inner and outer leaflets may be caused not only by PS but also by phosphatidylinositol 4,5-bisphosphate (PIP2), which is abundant in the inner leaflet of the plasma membrane of the cleavage furrow and epithelial AJCs. PIP2 promotes polymerization of the actin cytoskeleton and activation of membrane-anchored proteins such as ERMs, which may promote further cholesterol accumulation at the outer leaflet via stabilization of the actin cytoskeleton; it is also conceivable that the accumulation of cholesterol in the outer layer of the plasma membrane is promoted by proteins recruited by ERM. It is also notable that ZO proteins were essential for cholesterol accumulation at AJCs since it was reported that ZO-1 also accumulates at the cleavage furrow during cytokinesis (Wang et al., 2014). Therefore, it is possible that ZO proteins directly or indirectly regulate cholesterol accumulation to these membrane domains.

### Concluding remarks

It was long thought that ZO proteins bound to the actin cytoskeleton determine where claudins are assembled into functional TJs based on the analogy to other cell adhesion complexes such as AJs and desmosomes. It is increasingly clear, however, that TJs are fundamentally different structures. We previously demonstrated that removal of cholesterol from the plasma membrane selectively impairs TJ formation without affecting AJ formation (Shigetomi et al., 2018). Now, the present study shows that the scaffolding proteins and cytoskeleton associated with TJs promote the assembly of a functional adhesion structure by a mechanism completely unique from those of other cell adhesion structures. Altogether, our results strongly argue for a reconsideration of TJ formation, whereby protein-lipid (claudin-cholesterol) interaction—not protein-protein (claudin-ZO proteins) interaction as canonically believed—informs claudin strand polymerization.

## Materials and methods

### Cells and reagents

EpH4 cells were grown in DMEM supplemented with 10% FBS. Claudin-null EpH4 cells were established using the CRISPR-Cpf1 system. The primary and secondary antibodies used for immunofluorescence microscopy and immunoblotting are listed in Supplementary Table. Cytochalasin D was added to the cells at a final concentration of 1 µM.

### Fluorescence microscopy

Immunofluorescence microscopy was performed as described previously (Ikenouchi et al., 2005). In brief, cells cultured on coverslips were fixed with 3% formalin in PBS for 10 min at room temperature (RT), treated with 0.2% Triton X-100 in PBS for 5 min, and then washed with PBS. Blocking was done by incubating the fixed cells with 5% BSA in PBS for 30 min at RT. After the antibodies were diluted with the blocking solution, the cells were incubated at RT for 1 h with the primary antibody and for 1 h with the secondary antibody. For actin staining, Alexa Fluor 488 phalloidin (Life Technologies) was added to the secondary antibody. Specimens were observed at RT with a confocal microscope (LSM700; Carl Zeiss MicroImaging, Inc.)

Cells cultured on glass bottom dish. Then, cells incubated with recombinant GFP- or RFP-tagged D4 for 30 min at 37°C. After washed with HBSS, 3 times, cells were observed at 37°C with a confocal microscope (LSM700; Carl Zeiss MicroImaging, Inc.) equipped with a Plan-APO (63/1.40 NA oil-immersion) objective with appropriate binning of pixels and exposure time. The images were analyzed with ZEN 2012 (Carl Zeiss MicroImaging, Inc.).

### Freeze-fracture EM

Freeze fracture electron microscopy was performed as described previously (Shigetomi et al., 2018). Confluent cells were fixed with 2.5% glutaraldehyde in phosphate buffer, rinsed in phosphate buffer, cryoprotected in 30% glycerol in phosphate buffer, and then frozen in liquid propane. Frozen samples were fractured at −180 °C and platinum-shadowed unidirectionally at an angle of 45° in a JFD-7000 apparatus (JEOL). Replicas with cells were immersed in household bleach and then mounted on copper grids after the cells were removed. Processed samples were examined using a JEM-2100HCKM electron microscope.

### Measurement of Transepithelial Electric Resistance (TER) and Analysis of Paracellular Tracer Flux

Cells were cultured for six days on transwell filters and TER was measured directly in culture media using a cellZscopeE (CellSeed). For the paracellular tracer flux assay, after 6 days of culture, FITC-dextran with molecular weights of 3-5 kDa and 150 kDa at concentrations of 200 µM and 50 µM was added to the medium in the inner compartment. After 90 min of incubation, a 100 µl aliquot of the medium was collected from the outer compartment, and the paracellular tracer flux was measured as the amount of FITC-dextran in the medium using a fluorometer.

### Biotin tracer assay

The biotin tracer assay was performed using the cell-surface biotinylation method as described by (Chen et al., 1997) with some modifications. Cells cultured on transwell filters were washed with HEPES-buffered saline (HBS; 25 mM HEPES-NaOH [pH 7.2], 137 mM NaCl, 5 mM KCl, 0.7 mM Na2HPO4, 6 mM dextrose, and 1.8 mM CaCl2) to which 1 mg/ml EZ-Link sulfo-NHS-LC-biotin (Pierce) prepared in HBS was added to the upper chamber only. After 10 min incubation, cells were washed with DMEM and fixed with 100% methanol for 3 min at −20°C. Cells were then washed with PBS, blocked with 1% BSA in PBS for 30 min, and incubated with rat anti-E-cadherin antibody for 1 hr. After a wash with PBS, they were incubated for 30 min with streptavidin and anti-rat antibody conjugated with Texas Red and Cy2, respectively.

## Figure legends

**Supplementary Figure 1.**
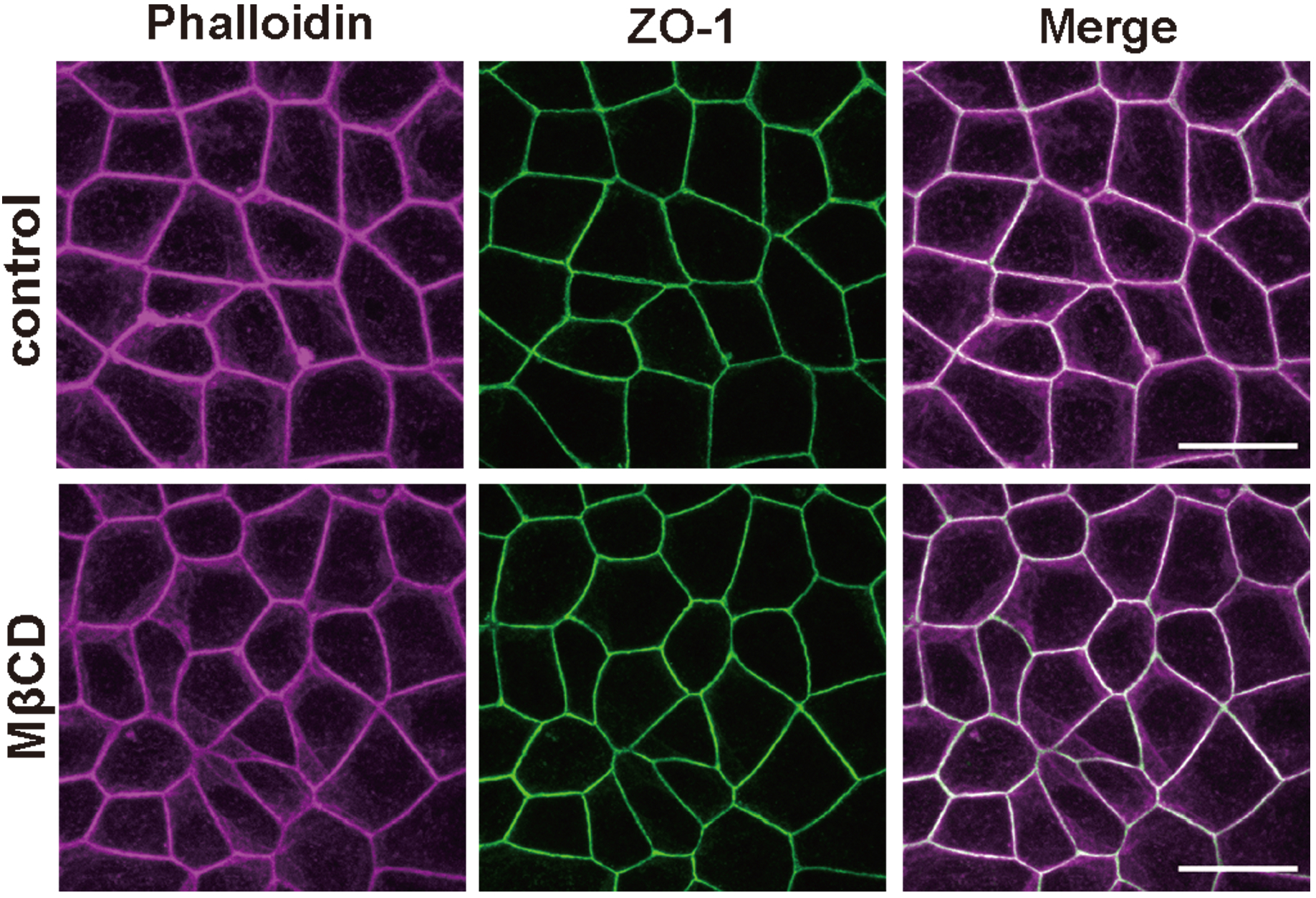
Removal of cholesterol from plasma membrane does not impair circumferential actin ring formation. Representative immunofluorescence images of MbCD-treated cells stained with phallidin and anti-ZO1 mAb. Scarle bar, Scale bar, 20 μm.

## Acknowledgements

We thank all members of the Ikenouchi laboratory (Department of Biology, Faculty of Sciences, Kyushu University, Fukuoka, Japan) for helpful discussions, and The Ultramicroscopy Research Center Kyushu University, Dr. T. Inai, K.Morishita (Fukuoka Dental College, Fukuoka, Japan), Dr. Y. Fukazawa and T. Maegawa (Fukui university, Fukui, Japan) for support of electron microscopy.

This work was supported by JSPS KAKENHI (JP19H03227 (J.I.), JP21K19231(J.I.), JP17J00211 (K.S.), and JP20K22632 (K.S.)), AMED-FORCE (21444781) (J.I.), JST-FOREST (JPMJFR204L) (J.I.) and grants from the Mitsubishi Foundation (J.I.) and the Cell Science Research Foundation (J.I.). The authors declare no competing financial interests.

## Author contributions

K. S. performed most of the experiments, analyzed the data and wrote the paper. J. I. designed the research and wrote the paper.

---

**supplementary Table.**
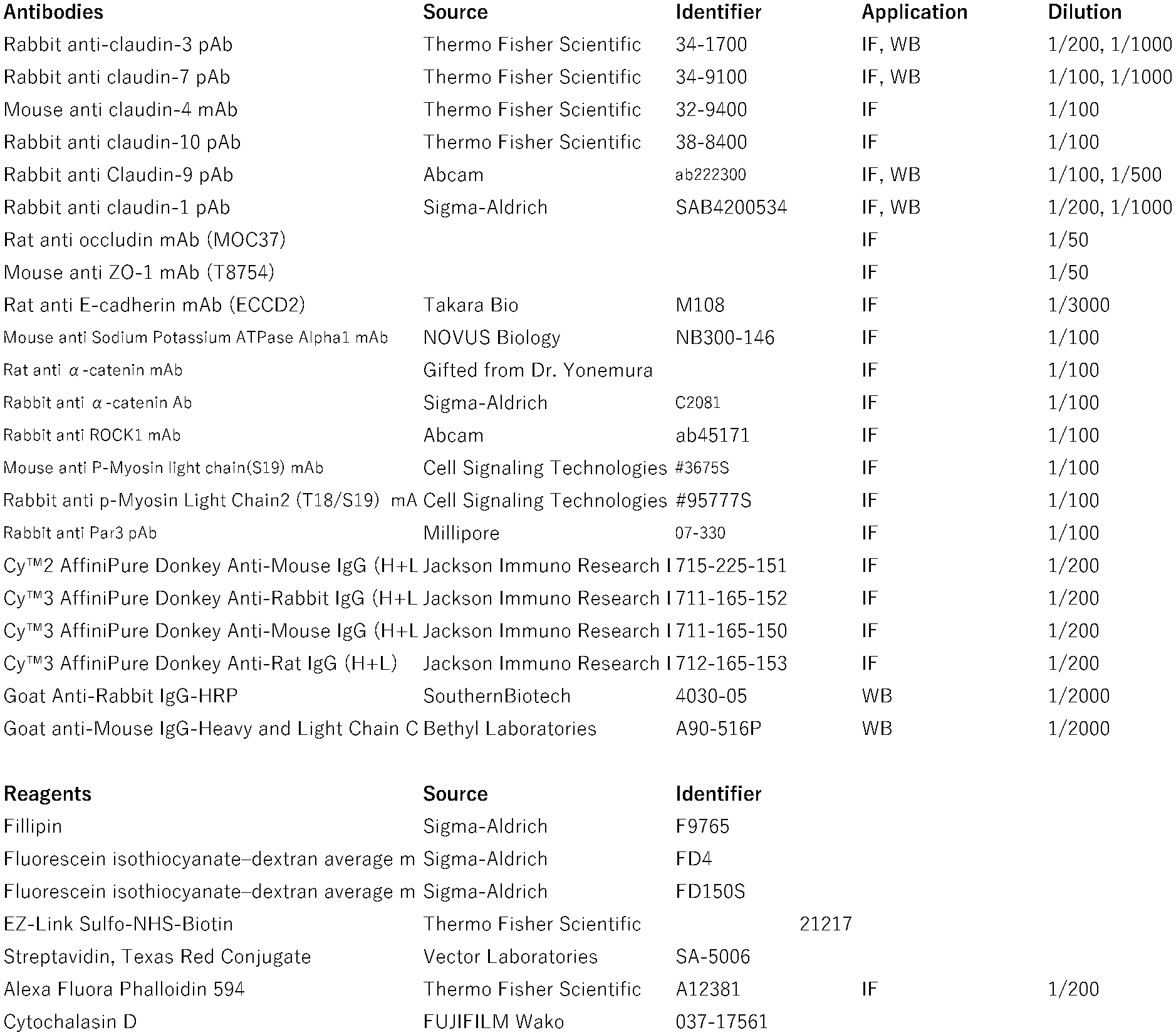

## Source Data

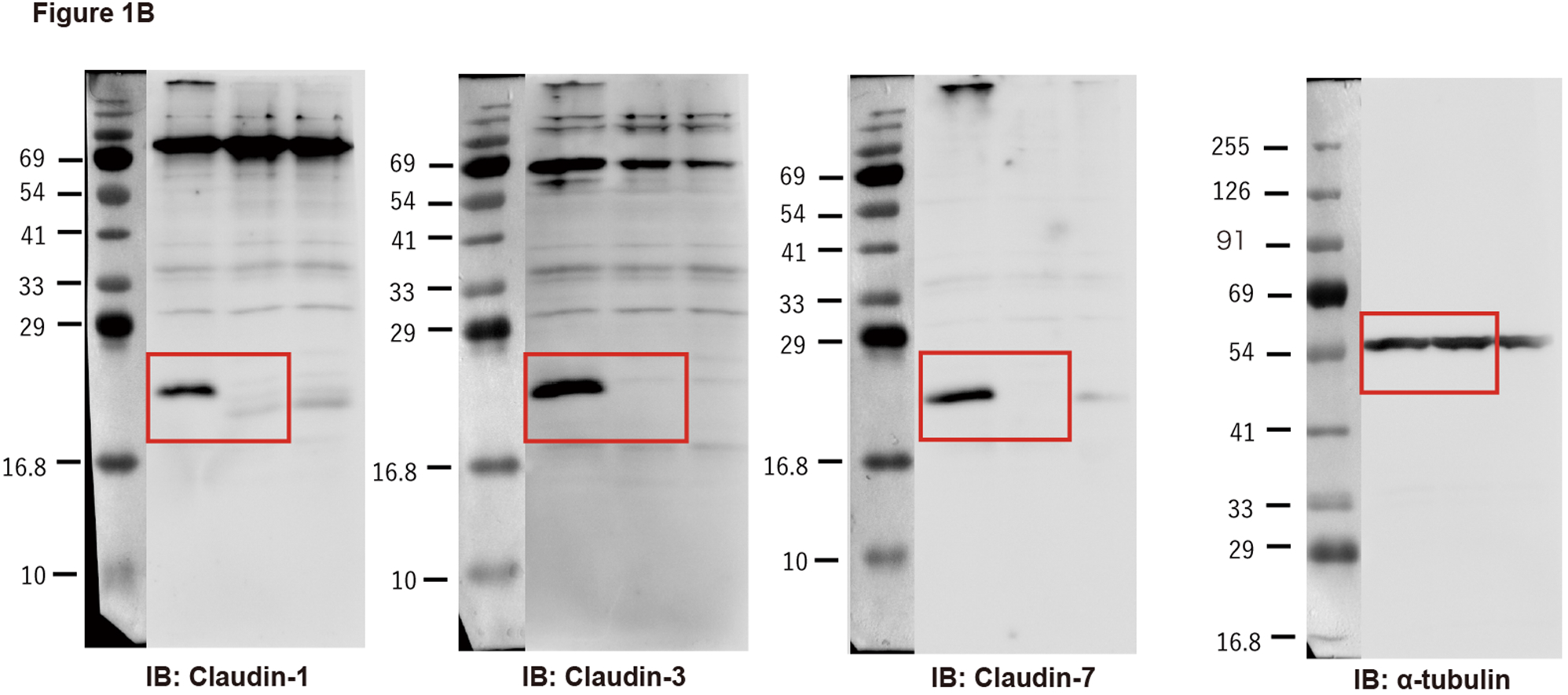

